# Inactivation of necroptosis-promoting protein MLKL creates a therapeutic vulnerability in colorectal cancer cells

**DOI:** 10.1101/2024.09.05.611491

**Authors:** Peijia Jiang, Sandhya Chipurupalli, Byong Hoon Yoo, Xiaoyang Liu, Kirill V. Rosen

**Author notes:** Correspondence: Dr. Kirill Rosen, Atlantic Research Centre, Rm C-304, CRC, 5849 University Avenue, PO Box 15000, Halifax, NS, B3H 4R2 Canada.

## Abstract

Mortality from colorectal cancer (CRC) is significant, and novel CRC therapies are needed. A pseudokinase MLKL typically executes necroptotic cell death, and MLKL inactivation protects cells from such death. However, we found unexpectedly that MLKL gene knockout enhanced CRC cell death caused by a protein synthesis inhibitor homoharringtonine used for chronic myeloid leukemia treatment. In an effort to explain this finding, we observed that MLKL gene knockout reduced CRC cell autophagy and rendered such autophagy critically dependent on the presence of VPS37A, a component of the ESCRT-I complex. Moreover, homoharringtonine-induced activation of p38 MAP kinase (p38MAPK) prevented VPS37A from supporting autophagy in MLKL-deficient cells and triggered their parthanatos, a cell death type driven by poly(ADP-ribose) polymerase hyperactivation. Finally, a pharmacological MLKL inhibitor necrosulfonamide strongly cooperated with homoharringtonine in suppressing CRC cell tumorigenicity in mice. Thus, while MLKL mediates necroptosis, MLKL protects CRC cells from death caused by drugs blocking basal autophagy, e.g., homoharringtonine, and MLKL inhibition creates a therapeutic vulnerability that could be utilized for CRC treatment.

## Introduction

20% of patients with colorectal cancer (CRC) have metastatic disease at the time of diagnosis, and 40% develop metastases after treatment of the localized disease^1^. 5-year survival of patients with metastatic CRC is below 20%^1^. Hence, novel CRC therapies are needed.

Necroptosis is programmed necrosis driven by protein kinases RIPK1 and RIPK3^2^. When activated by various stimuli, RIPK1 binds RIPK3, which is then activated by autophosphorylation. RIPK3 further phosphorylates and thereby activates a pseudokinase MLKL^2^. MLKL then polymerizes, forms pores in the plasma membrane and thus kills the cell^2^.

We noticed that a well-established and highly specific small molecule MLKL inhibitor necrosulfonamide^3^ or MLKL gene knockout (KO) reduce CRC cell survival. This finding was unexpected since MLKL is well known to promote cell death during necroptosis, rather than protect the cells from death^2^. Nevertheless, since MLKL gene KO is non-toxic to mice^4^ but seems to be toxic to CRC cells, we reasoned that MLKL inactivation might represent a novel approach for CRC treatment.

We found that CRC cell death caused by MLKL inactivation is enhanced by homoharringtonine (HHT), an alkaloid, also known as omacetaxine, currently used for treatment of chronic myelogenous leukemia^5^. We found that MLKL inactivation and HHT cooperate with each other in blocking ESCRT machinery-dependent basal autophagy of CRC cells and kill them by a form of cell death termed parthanatos.

HHT is thought to kill cells by inhibiting the activity of the ribosomes^6^. Similar to other ribosome inhibitors, HHT triggers ribotoxic stress response, a chain of events that starts with inactivation of the ribosome and then triggers activation of protein kinases p38 MAP kinase (p38MAPK) and JNK which either allows the cell to survive through the stress or kills the cell if the stress magnitude or duration is incompatible with cell survival^7^.

Autophagy is a process of degradation of the cellular content mediated by the double-membrane vesicles termed autophagosomes^2^. One of the functions of autophagy is to promote cell survival by eliminating the toxic cellular content, including damaged proteins and organelles^8^. In addition, when cells experience lack of nutrients, autophagy rescues cells from death by promoting degradation of the non-essential cellular components to provide the building material for the essential ones^9^.

Endosomal sorting complexes required for transport (ESCRTs) mediate the formation of numerous types of vesicles, including endosomes, autophagosomes and extracellular vesicles^10^. ESCRT machinery consists of the ESCRT-I, -II and -III complexes and was found to promote autophagy via several mechanisms^11, 12^.

Parthanatos is a form of programmed necrosis triggered by hyperactivation of poly(ADP-ribose) polymerase 1 (PARP1), an enzyme that promotes the formation of poly(ADP-ribose) (PAR) in the cell^13^. When produced at excessive levels, PAR binds a cell death-promoting factor AIF localized to the mitochondrial outer membrane. This causes the release of AIF from the mitochondria. AIF further binds an endonuclease MIF, the AIF–MIF complex translocates to the nucleus where it causes chromosomal DNA degradation and cell death^13^. Notably, PARP1 is overexpressed in human CRC cells compared to normal intestinal epithelial cells and promotes colon cancer in mice^14^. Thus, high level and activity of PARP1 in CRC cells could make them vulnerable to PARP1-hyperactivating therapies and thereby create a therapeutic window for CRC treatment with parthanatos-inducing drugs.

We found here that MLKL inactivation renders CRC cell autophagy strongly dependent on the presence of the ESCRT-I complex component VPS37A^10^ and that in the absence of MLKL, HHT activates p38MAPK and thereby prevents VPS37A from supporting autophagy. We observed that as a result of autophagy inhibition caused by the combined effect of MLKL inactivation and HHT, CRC cells die by parthanatos. Moreover, we found that an MLKL inhibitor necrosulfonamide and HHT strongly cooperate with each other in suppressing the ability of CRC cells to form tumors in mice. Hence, pharmacological MLKL inhibition combined with HHT treatment represents a potential novel approach for CRC therapy.

## Results

### MLKL inactivation reduces clonogenic survival of CRC cells

Activating mutations of RAS GTPase often occur in CRC^15^ and drive CRC progression^16^. In the course of our earlier studies of the mechanisms of RAS-dependent CRC cell survival we found that RAS-induced activation of a protein kinase ERK promotes CRC cell survival by blocking both apoptotic and non-apoptotic cell death signals^17^. The non-apoptotic events blocked by the RAS/ERK signaling pathway in CRC are not well understood and we decided to explore them.

Necroptosis, a form of non-apoptotic programmed cell death driven by plasma membrane permeabilization, occurs in response to stimuli, such as viral infection^18^. A pseudokinase MLKL, a key necroptosis mediator, drives cell death by forming an oligomeric structure that disrupts the cell membrane^19^. To test whether ERK inhibition in CRC cells causes necroptosis we examined whether death of oncogenic RAS-carrying CRC cells HCT116^20^ caused by the ERK inhibitor SCH 772984^21^ is blocked by necrosulfonamide (NSA), a widely used small molecule MLKL inhibitor that binds MLKL at Cys86^3^ and blocks MLKL-driven necroptosis by preventing MLKL incorporation in the cell membrane^3^. To assess cell viability, we used clonogenic survival assay often utilized for this puropose^17, 22^. We found that NSA did not block SCH 772984-induced cell death (Fig. 1a) indicating that inhibition of necroptosis is not the mechanism by which ERK promotes CRC cell survival.

**Fig. 1.**
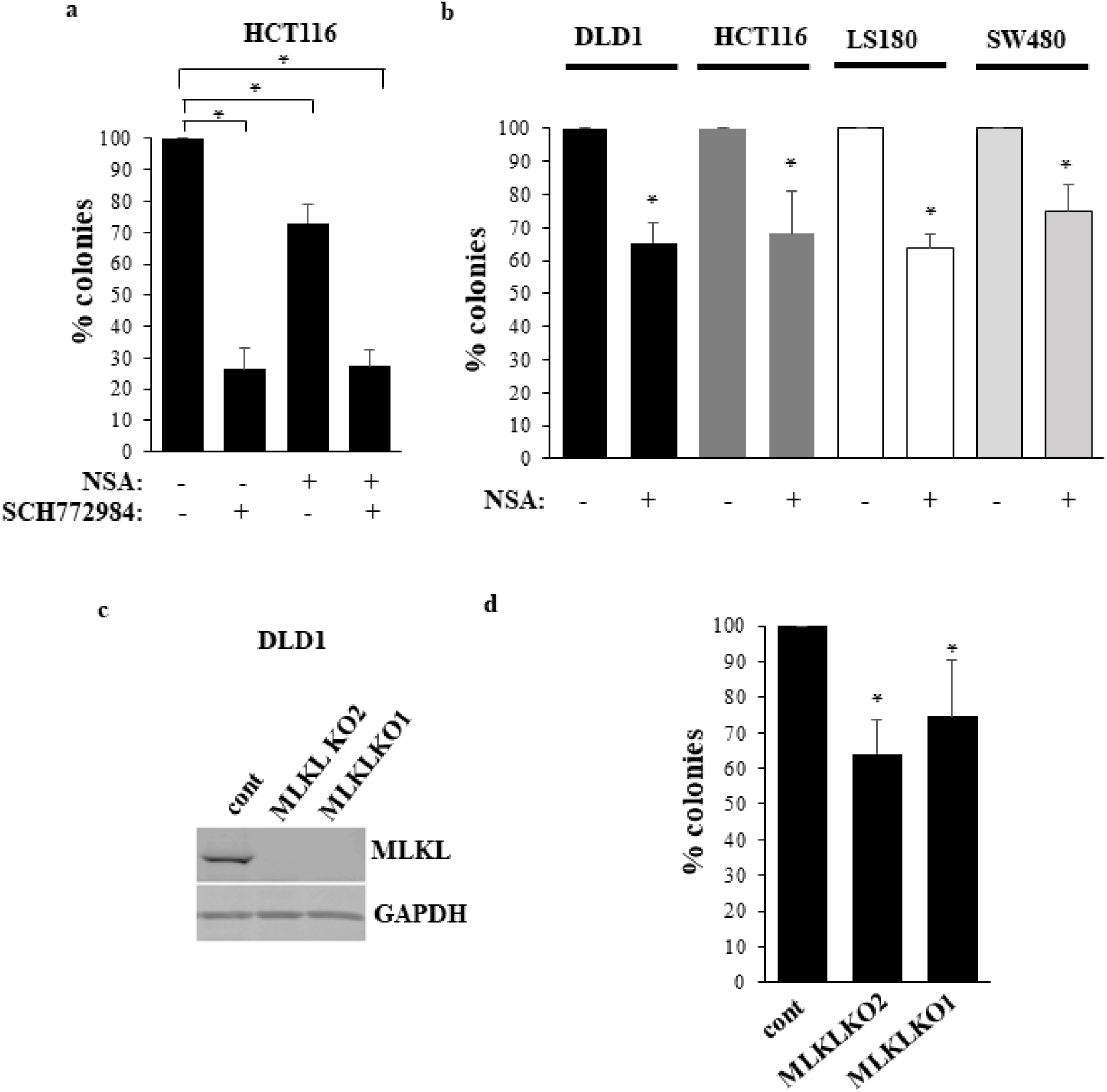
MLKL inactivation reduces CRC cell survival. **a** HCT116 cells were treated with DMSO (-) or 1μM SCH 772984 (SCH) and/or 3 μM NSA (+), and colonies formed by the cells were counted. **b** Indicated cell lines were treated with DMSO (-) or 1 μM NSA (+), and colonies formed by the cells were counted. **c** Control cells (cont) or MLKL-deficient cells MLKL KO1 and MLKL KO2 were tested for MLKL levels by western blotting. GAPDH served as a loading control. **d** Cells generated in **c** were assayed as in **a**. The data in **a** are the average of the triplicates plus SD, and this experiment was repeated twice with similar results. The data in **b** and **d** are the average of three independent experiments plus SD. * p value < 0.05.

Unexpectedly, we also noticed that NSA caused moderate but noticeable reduction of the clonogenic survival of HCT116 cells (Fig. 1a). This finding suggests that MLKL promotes cell survival, even though MLKL is well known to drive cell death during necroptosis^18^. We were intrigued by this observation and focused further studies on testing whether MLKL can indeed protect cells from death and whether this MLKL property can potentially serve for CRC treatment. To this end, we first tested whether NSA inhibits survival of other CRC cell lines. Similarly to what we observed for HCT116 cells, we found that NSA moderately reduced clonogenic survival of CRC cells LS180^17^, DLD1^23^ and SW480^24^ (Fig. 1b). To verify these data by a genetic approach we transfected DLD1, one of these cell lines, with a CRISPR/Cas9 vector encoding single guide (sg)RNAs targeting MLKL exons 2 or 9 and generated two cell clones in each of which MLKL gene was disrupted by one of these sgRNAs (Fig. 1c). We found that MLKL-deficient cells are noticeably less clonogenic than the control, MLKL-expressing cells (Fig. 1d). Thus, MLKL inactivation reduces clonogenic survival of CRC cells.

### HHT enhances CRC cell death caused by MLKL inactivation

Remarkably, *MLKL*-deficient mice are viable, healthy, fertile, are born with normal Mendelian frequency and do not show any physical or behavioral abnormalities^4^. Since MLKL inactivation is non-toxic to mice^4^ but seems to be toxic to CRC cells (Fig. 1a, b, d), we reasoned that, if enhanced further, death signals caused by such inactivation could be used for CRC treatment. To this end, we tested whether drugs targeting various CRC-promoting signaling pathways, e.g. vemurafenib (a RAF inhibitor)^25^, selumetinib (a MEK inhibitor)^26^, pictilisib (a PI3 kinase inhibitor)^27^ or lapatinib (an EGFR/ERBB2 inhibitor)^28^, enhance death of DLD1 cells caused by MLKL gene knockout (KO). However, none of the drugs exerted this effect (Fig. 2a). We further noticed that HHT, an alkaloid, also known as omacetaxine, currently used for treatment of chronic myelogenous leukemia^5^ and is thought to kill cells by inhibiting the ribosomes^6^, strongly cooperated with MLKL gene deletion in reducing clonogenic survival of DLD1 cells (Fig. 2a). We observed that HHT reduces not only clonogenicity but also the total number of MLKL-deficient DLD1 cells significantly more effectively than that of the control MLKL-expressing cells (Fig. 2b). We also found that HHT and MLKL inhibitor NSA noticeably cooperate with each other in inhibiting clonogenic survival of CRC cell lines DLD1, HCT116, LS180 and SW480 (Fig. 2c-f). Thus, genetic or pharmacological MLKL inhibition cooperates with HHT in blocking CRC cell survival.

**Fig. 2.**
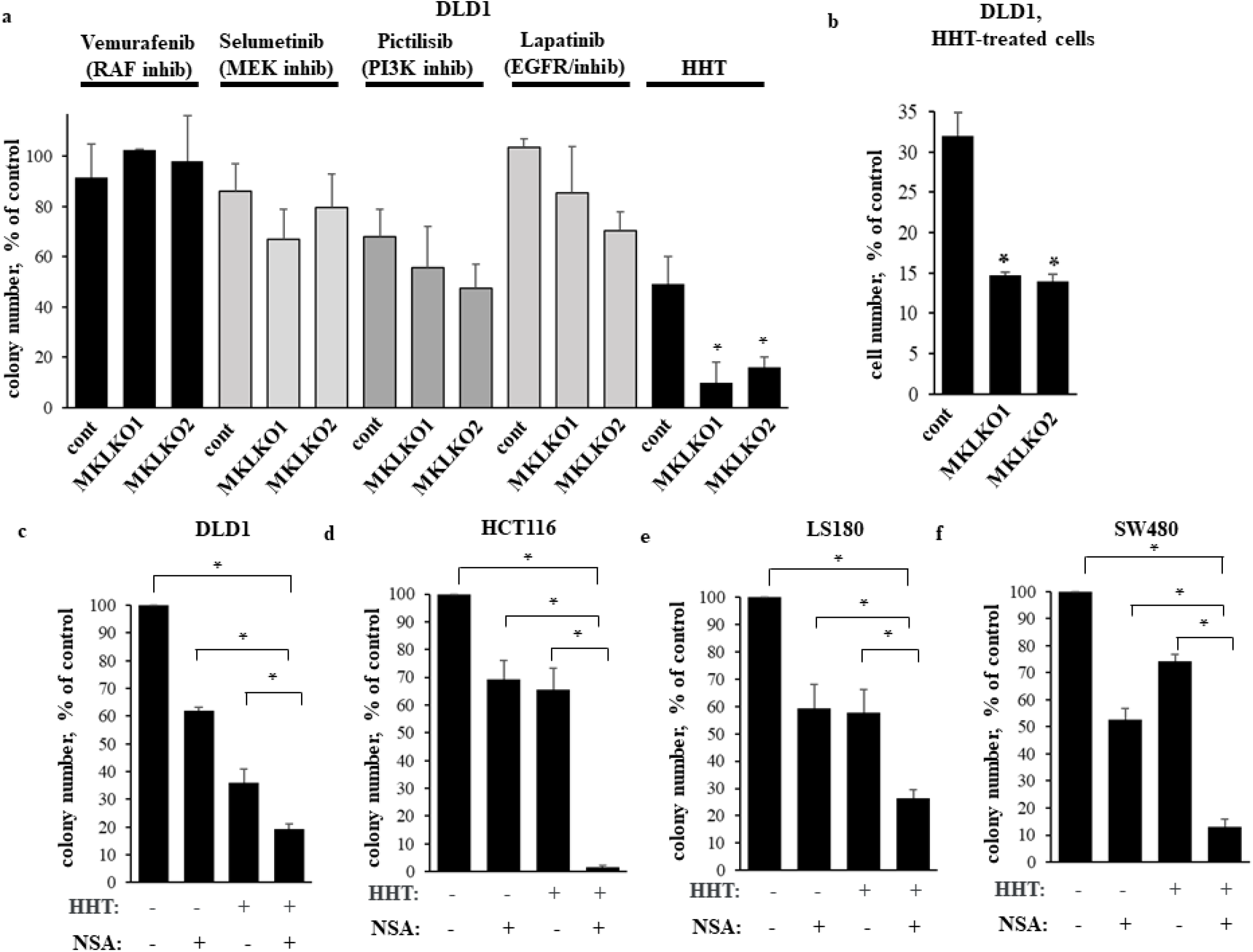
MLKL inactivation cooperates with HHT in reducing CRC cell survival. **a** Indicated cell lines were treated or not with 5μM vemurafrnib, 1 μM selumetinib, 1 μM pictilisib, 1 μM lapatinib or 10 ng/ml HHT and colonies formed by the cells were counted**. b** Indicated cell lines were treated or not with 10 ng/ml HHT for 96h and counted**. c-f** Indicated cell lines were treated (+) or not (-) with 1 μM NSA and 10 ng/ml HHT, and colonies formed by the cells were counted. The data represent % of colony number formed by the untreated cells (**a**, **c-f**) or % of untreatded cells (**b**). The data in **a** are the average of two independent experiments plus SD, those in **c-f**, the average of the triplicates plus SD, and these expriments were repeated twice with similar results. * p value < 0.05.

### NSA and HHT cooperate in reducing CRC cell tumorigenicity *in vivo*

Given that NSA and HHT cooperated in reducing clonogenic survival of multiple CRC cell lines in culture, we tested whether these drugs cooperate in suppressing tumorigenicity of these cells *in vivo*. To this end, we subcutaneously injected one of these cell lines, HCT116, in the immunodeficient mice, allowed the tumors to form and then injected the mice with DMSO (vehicle), NSA, HHT or both drugs. As shown in Fig. 3a the two drugs noticeably cooperated in reducing growth of respective tumors. Of note, the drug combination did not seem to be toxic to the mice as they did not lose weight in response to treatment with each drug alone or both together (Fig. 3b) and did not display any gross physical abnormalities. Thus, NSA and HHT significantly cooperate with each other in suppressing tumorigenicity of CRC cells *in vivo*.

**Fig. 3.**
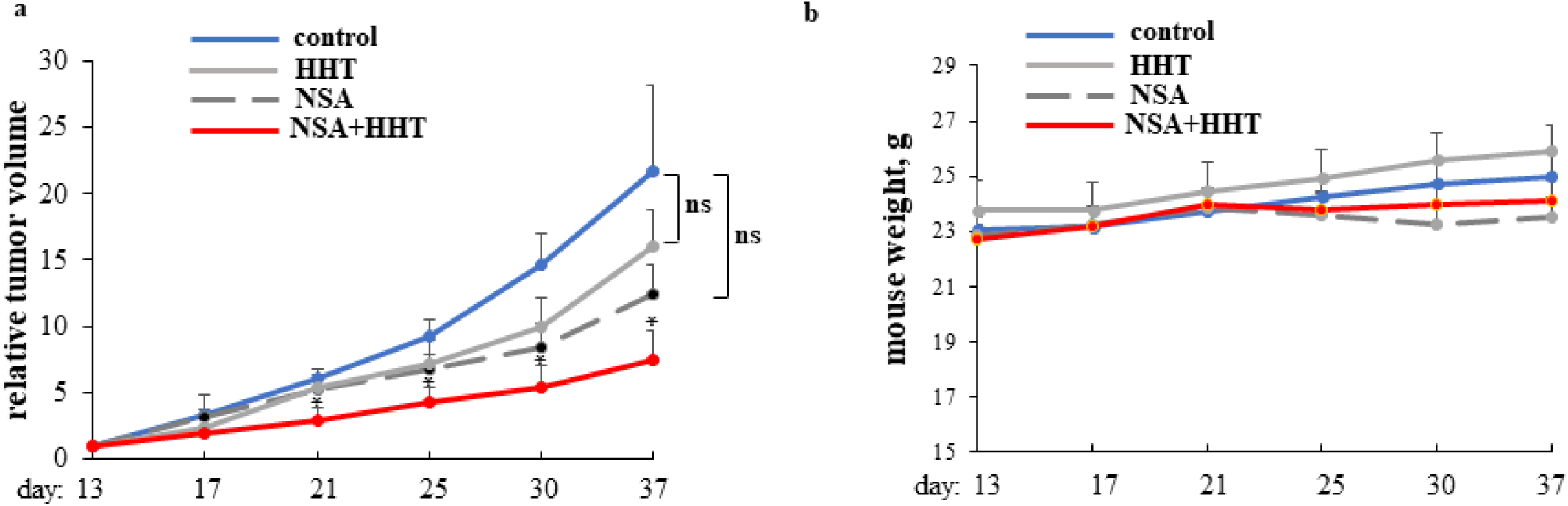
MLKL inhibitor NSA cooperates with HHT in blocking CRC cell tumorigenicity *in vivo*. **a** HCT116 cells were injected in the flanks of 28 Nu/Nu Nude mice. Once tumor volumes reached the average volume of 80 mm^3^, the mice were injected intraperitoneally with DMSO (control), NSA, HHT or with NSA and HHT together. 7 mice were used per group. Changes in relative tumor volumes **a** and mouse weight **b** plus SE are shown. Tumor volume observed on the day of the first injection was designated as 1.0 for each mouse, and subsequent changes in each tumor volume were calculated relative to that number. *- p ˂0.05.

### MLKL does not protect CRC cells from death by secreting soluble factors or promoting TRAIL receptor degradation

Given that MLKL inactivation combined with HHT blocks CRC cell growth *in vitro* and *in vivo* (Fig. 2, 3), and, therefore, represents potential novel approach for CRC therapy, we felt that it would be of interest to understand molecular mechanisms by which this combination treatment kills CRC cell. Elements of such mechanisms could serve as future targets for or biomarkers of CRC sensitivity to such therapies. Interestingly, MLKL was suggested to trigger pro-survival signals in certain circumstances, and we tested whether these circumstances apply to our model.

It was proposed that MLKL drives secretion of the pro-survival cytokines by breast tumor cells and that these cytokines induce cellular paracrine survival signals^29^. This notion is based on the data showing that cell growth blocked by MLKL gene knockout can be restored by the conditioned medium derived from the parental MLKL-expressing cells^29^. However, we found that MLKL-deficient DLD1 cells cannot be rescued from HHT treatment by the conditioned medium derived from the MLKL-producing control cells (Fig. 4a).

**Fig. 4.**
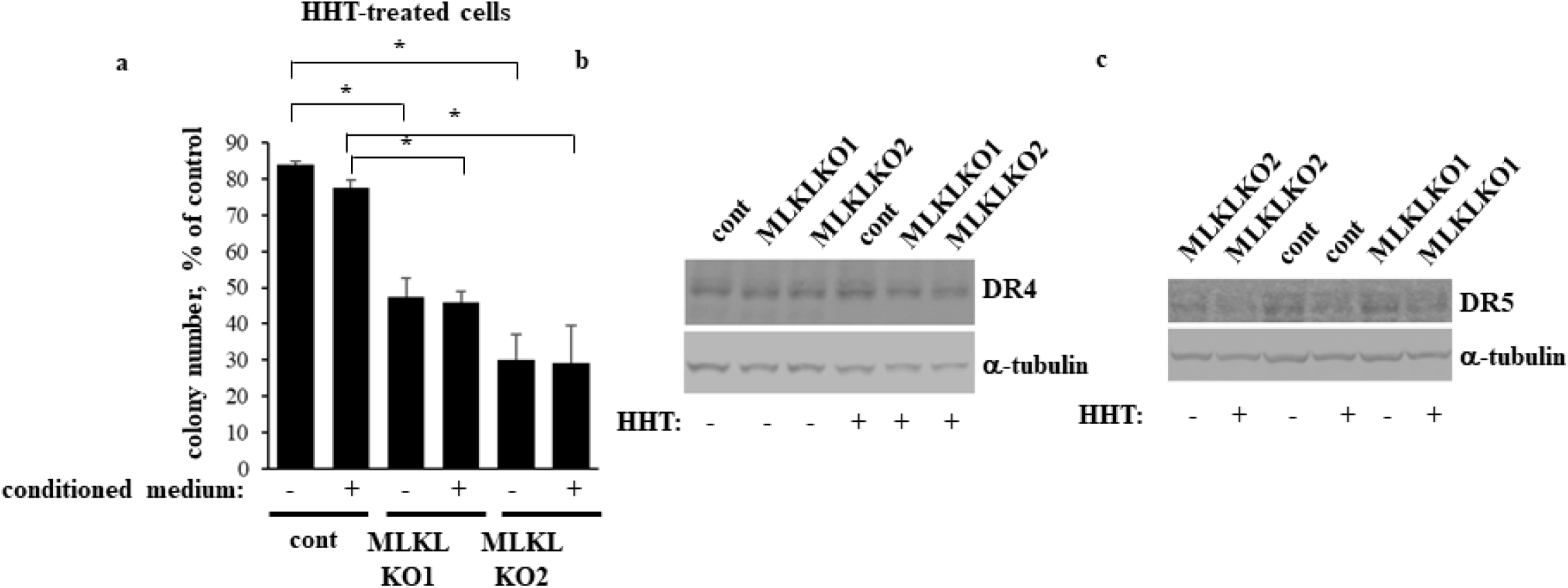
MLKL does not protect CRC cells from death by secreting soluble factors or promoting TRAIL receptor degradation. **a** Indicated cell lines were treated (+) or not (-) with 10 ng/ml HHT in the absence (-) or in the presence (+) of the conditioned medium derived from the same number of control untreated cells, and colonies formed by the cells were counted. The data in **a** are the average of three independent experiments plus SD. * p value < 0.05. **b, c** Indicated cell lines were treated as in **a** for 72h **a** or 48h **b** and assayed for DR4 (**a**) or DR5 (**b**) levels by western blotting.

MLKL gene knockdown was also shown to block lysosomal degradation of the death receptor DR5 in lung and breast tumor cells by unknown mechanisms and thereby enhance cell death upon treatment with exogenous TRAIL, the ligand for this receptor^30^. Since CRC cell lines do produce TRAIL^31, 32^, it is conceivable that if DR5 is upregulated in MLKL-deficient CRC cells, this receptor could sensitize the cells to the endogenously produced TRAIL, either by itself or in the presence of HHT. However, we found that MLKL loss upregulates neither DR5 nor DR4, another TRAIL receptor^2^, in CRC cells neither before nor after HHT treatment (Fig. 4b, c). Thus, none of the indicated mechanisms seem to mediate the effect of MLKL on CRC cells observed by us.

### MLKL inactivation and HHT cooperate in killing CRC cells by inhibiting their basal autophagy

Since autophagy is a well-known regulator of cell survival^2^, we investigated whether HHT and NSA cooperate with each other in blocking the basal autophagy of CRC cells. Lipidation of autophagy-driving protein LC3B is a major mechanism of autophagy, and formation of the lipidated LC3B form termed LC3B-II is a well-established autophagy marker^33^. Of note, LC3B-II is eventually degraded during autophagy after the fusion of the autophagosomes to the lysosomes. Hence, while LC3B-II upregulation can signify increased autophagy, it can also be the symptom of autophagy inhibition caused by reduced autophagosome-to-lysosome fusion^34^. To ensure that increased LC3B-II levels reflect increased autophagy in the studies outlined below, we tested the effect of HHT and NSA on LC3B-II in the absence and in the presence of bafilomycin A1, a lysosomal inhibitor that disrupts the autophagic flux by blocking the fusion of the autophagosomes to the lysosomes and thereby prevents the resulting LC3B-II degradation^34^.

We tested whether MLKL KO or pharmacological MLKL inhibition cooperates with HHT in reducing LC3 lipidation in DLD1 cells. LC3B-II was poorly detectable in bafilomycin A1-untreated cells, while bafilomycin A1 noticeably upregulated LC3B-II in DLD1 cells and their MLKL KO variants (Fig. 5a, b). These data indicate that the basal rate of autophagy in the cells is sufficiently high to rapidly trigger LC3B-II lysosomal degradation and thereby render LC3B-II poorly detectable, unless autophagosome-to-lysosome fusion is blocked. We further observed that MLKL KO (Fig. 5a, b), NSA (Fig. 5c) or HHT (Fig. 5a-c) noticeably downregulated LC3B-II in bafilomycin A1-treated MLKL-expressing DLD1 cells. Remarkably, MLKL KD or NSA treatment significantly cooperated with HHT in downregulating LC3B-II (Fig. 5a-c). Thus, MLKL inactivation and HHT cooperate in reducing the basal autophagy of CRC cells.

**Fig. 5.**
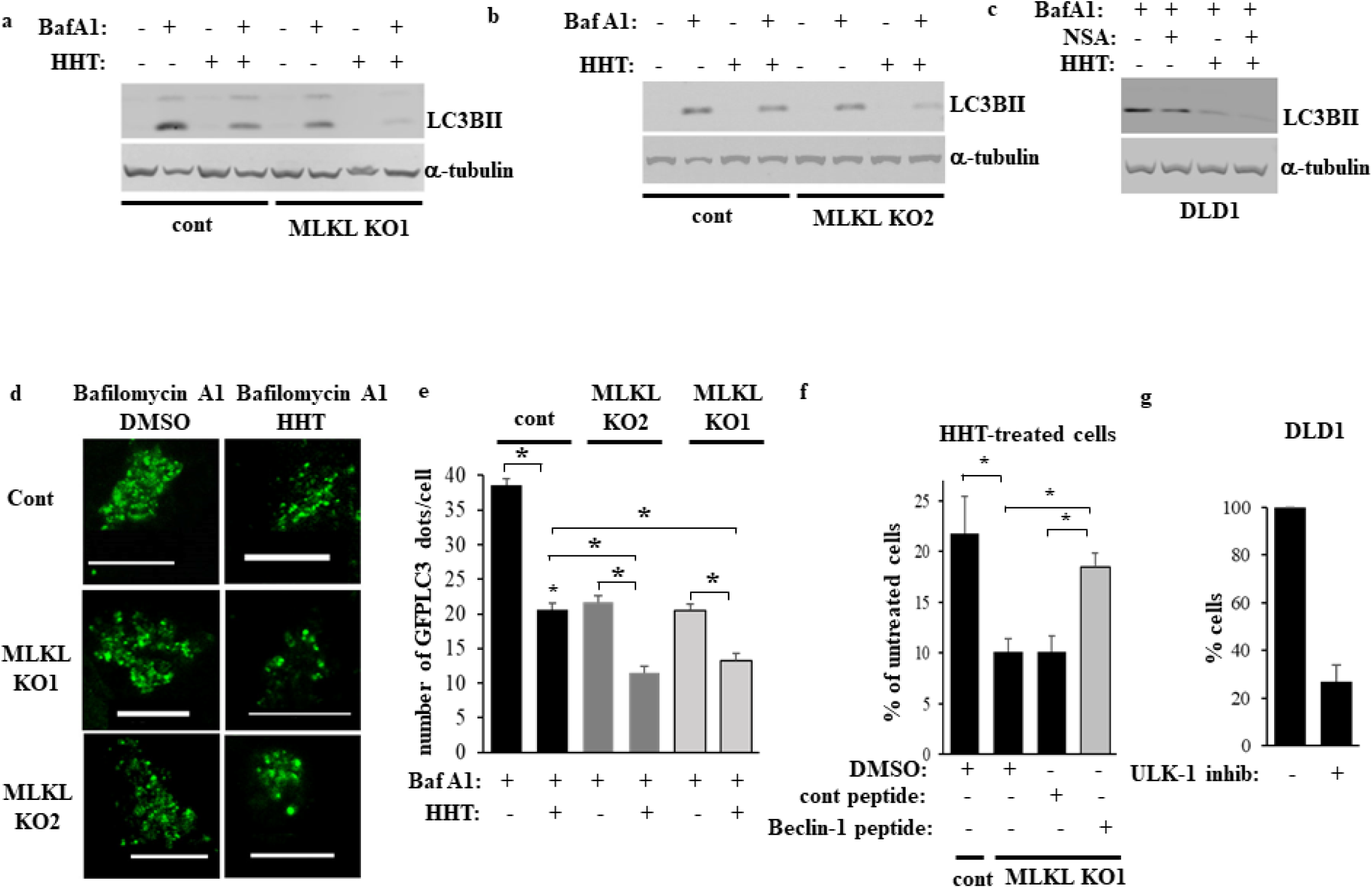
MLKL inactivation and HHT cooperate in killing CRC cells by inhibiting their basal autophagy. **a, b** Indicated cell lines were treated with DMSO (-) or 10 ng/ml HHT (+) for 24h in the absence (-) or in the presence (+) of 50 nM bafilomycin A1 and the cells were assayed for LC3B levels by western blotting. **c** Indicated cell lines were treated with DMSO (-) or 10 ng/ml HHT and/or 3 μM NSA (+) for 24h in the presence of 50 nM bafilomycin A1 and the cells were assayed for LC3B levels by western blotting. α-tubulin was used as a loading control in **a-c**. **d** Indicated cells were transfected with the GFP-LC3 expression vector and treated with DMSO (DMSO) or 10 ng/ml HHT (HHT) for 24 in the presence of 50 nM bafilomycin A1. Green puncta per cell were counted. Representative fluorescence microscopy images are shown. Bar - 10 μm. **e** Quantification of the number of green puncta per cell for the cells treated as in **a**. The numbers represent the average of the number of puncta per cell plus the SE. 24 untreated and 13 HHT-treated cells were counted in the case of the control clone, 42 untreated and 28 HHT-treated cells in the case of MLKL KO1 cells, and 29 untreated and 34 HHT-treated cells in the case of MLKL KO2 cells This experiment was repeated twice with similar results. **f** Indicated cell lines were treated with DMSO (-) or 10 ng/ml HHT (+) for 96h in the absence (-) or in the presence (+) of 12 μM of control (cont) or cell-permeable Beclin-1 (Beclin-1) peptide and counted. **g** DLD1 cells were treated with DMSO (-) or 20 μM ULK-1 inhibitor (ULK-1 inhib) SBI-0206965 (+) for 96h and counted. The data in (**f**, **g**) are the average of three independent experiments plus SD. *- p ˂0.05.

To confirm our findings by a complementary approach, we examined the ability of green-fluorescent protein (GFP)-tagged LC3B protein (GFPLC3) to cause puncta formation in the control and MLKL KO CRC cells before and after bafilomycin A1 and HHT treatment. Formation of these puncta is a widely used marker of autophagosome formation^34^. Similar to what we observed in the case of the LC3BII western blots, GFPLC3 puncta were poorly detectable in the cells not treated with bafilomycin A1 (not shown) transiently transfected with a GFPLC3-encoding expression vector but were readily detectable in bafilomycin A1-treated cells (Fig. 5d, e). We found MLKL KO and HHT treatment significantly cooperated in reducing the number of the GFPLC3 puncta per cell in the respective cell lines (Fig. 5d, e). These data further support our conclusion that MLKL inactivation and HHT cooperate with each other in reducing the basal autophagy of CRC cells.

To test whether autophagy inhibition triggered by HHT in MLKL-deficient cells contributes to CRC cell loss observed by us we treated the MLKL KO cells with HHT and a cell permeable peptide composed of the HIV-1 TAT protein transduction domain attached to either a scrambled amino acid sequence (a control peptide) or the peptide derived from a protein Beclin-1, a major autophagy inducer^35^. The TAT domain renders the peptides cell-permeable while the indicated sequence of Beclin 1 is necessary and sufficient for autophagy induction^35^. The latter peptide efficiently induces autophagy and is being widely used as an autophagy-promoting tool^35, 36, 37^. We noticed that as expected, MLKL KO cells were significantly more sensitive to HHT than the control MLKL-expressing cells and that the control peptide did not rescue the MLKL KO cells from HHT (Fig. 5f). In contrast, the Beclin-1-derived peptide almost completely restored the number of HHT treated MLKL KO cells compared to that of HHT-treated MLKL-expressing control cells (Fig. 5f). Thus, inhibition of autophagy caused by MLKL inactivation and HHT is required for CRC cell death induced by this combination treatment. In this regard, our findings that RAF, MEK, PI3K and EGFR inhibitors did not cooperate with MLKL in killing CRC cells are not surprising since these agents were demonstrated to promote, rather than inhibit, autophagy in various cancer cell types, including CRC cells^38, 39, 40^. We further observed that SBI-0206965, a highly specific inhibitor of the critical autophagy driver protein kinase ULK1^41^, strongly reduced growth of MLKL-expressing control DLD1 cells (Fig. 5f). Hence, inhibition of autophagy is sufficient for suppressing CRC cell growth.

### MLKL inactivation renders CRC cell autophagy VPS37A-dependent

The ESCRT machinery includes the ESCRT-I, -II and III multiprotein complexes that mediate numerus vesicle-dependent cellular events, including endosomal trafficking, nuclear membrane assembly and promote the formation of various vesicles, such as endosomes, autophagosomes and extracellular vesicles^10^. When others subjected cells to necroptosis-promoting stimuli, MLKL was found to trigger the activity of the ESCRT-I and III complexes which repaired the MLKL-permeabilized cell membrane and rescued the cells from necroptosis until the cells irreversibly committed to die^42^. Moreover, the ESCRT complexes, including ESCRT-I, were found to promote autophagy via multiple and only partly understood mechanisms^11, 12^. Collectively, these data prompted us to investigate the role of the ESCRT-I machinery in the effects observed by us. To this end, we studied LC3BII levels in bafilomycin A1-treated MLKL-expressing and MLKL KO DLD1 cells before and after knocking down (KD) the ESCRT-I complex component VPS37A, which is critical for the function of the complex^43^, using two different VPS37A-specific siRNAs (Fig. 6a-d). We noticed that VPS37A KD did not have any effect on LC3B lipidation of the control, MLKL-expressing DLD1 cells (Fig. 6e, f). However, LC3B lipidation was strongly reduced in the MLKL KO DLD1 cells, indicating that such lipidation becomes VPS37A-dependent in the absence of MLKL (Fig. 6e, f). Moreover, VPS37A was unable to promote/support LC3B lipidation in MLKL-deficient cells when they were treated with HHT (Fig. 6e, f). Hence, MLKL inactivation reduces CRC cell autophagy (Fig. 5, 6) and renders this autophagy dependent on the presence of VPS37A. The latter protein is unable to promote autophagy if the cells are treated with HHT (Fig. 6e, f).

**Fig. 6.**
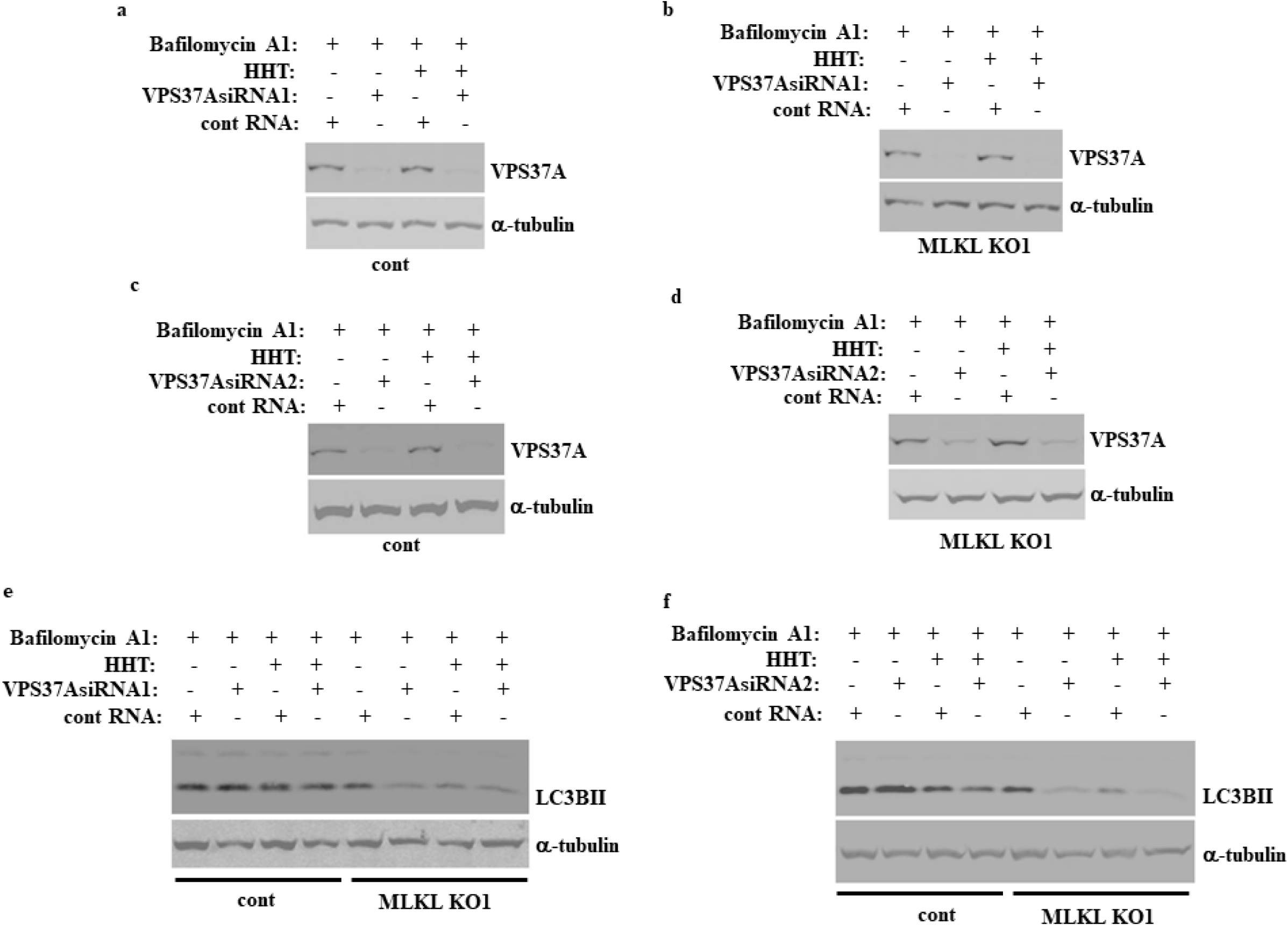
MLKL inactivation renders CRC cell autophagy VPS37A-dependent. Indicated cell lines were transfected with a 100 nM control RNA (cont RNA) or VPS37A-specidic siRNA (VPS37A siRNA) 1 (**a, b, e**) or 2 (**c, d, f**), treated with DMSO (-) or 10 ng/ml HHT (+) for 24h in the presence of 50 nM bafilomycin A1 and assayed for VPS37A (**a-d**) or LC3B (**e, f**) levels by western blotting. α-tubulin was used as a loading control.

### HHT contributes to autophagy inhibition in CRC cells by activating p38MAPK

Inhibitors of the ribosome function, including HHT, trigger ribotoxic stress response, a chain of signaling events that start with the ribosome inhibition and allow the cell to either survive the stress or kill the cell if the stress magnitude or duration is incompatible with cell survival^7^. The response was proposed to be triggered by a protein kinase ZAKα that is constitutively bound to translating ribosomes and is activated when the ribosomes are inhibited. Activated ZAKα phosphorylates and activates MAP kinases MKK3 and/or MKK6 which in turn phosphorylate and thereby activate p38MAPK and JNK^7^. This HHT-induced mechanism attracted our attention because p38MAPK is well known to inhibit autophagy via multiple signaling events, including phosphorylation and inactivation of the autophagy stimulator ATG5^44^ and sequestration of the p38-inetracting protein p38IP, whose binding to another autophagy stimulator ATG9 is required for autophagy execution^45^.

We found that HHT upregulates phospho-p38MAPK, a well-established sign of p38MAPK activation^46^, in both the control and MLKL KO DLD1 cells (Fig. 7a). Four members of the p38MAPK family are known, α, β, γ and δ^47^. We found that treatment of the control DLD1 cells and their MLKL KO variants with SB203580, a widely used specific small molecule p38MAPK α and β inhibitor^48^, in the presence of bafilomycin A1 upregulates LC3BII in all cases (Fig. 7a, b). Importantly, the inhibitor restored LC3BII levels in HHT-treated MLKL KO cells to the levels comparable to those observed in the case of untreated MLKL KO cells (Fig. 7b, c). Hence, HHT-induced p38MAPK activation is likely primarily responsible for the ability of HHT to cooperate with MLKL inhibition in blocking the basal autophagy in CRC cells.

**Fig. 7.**
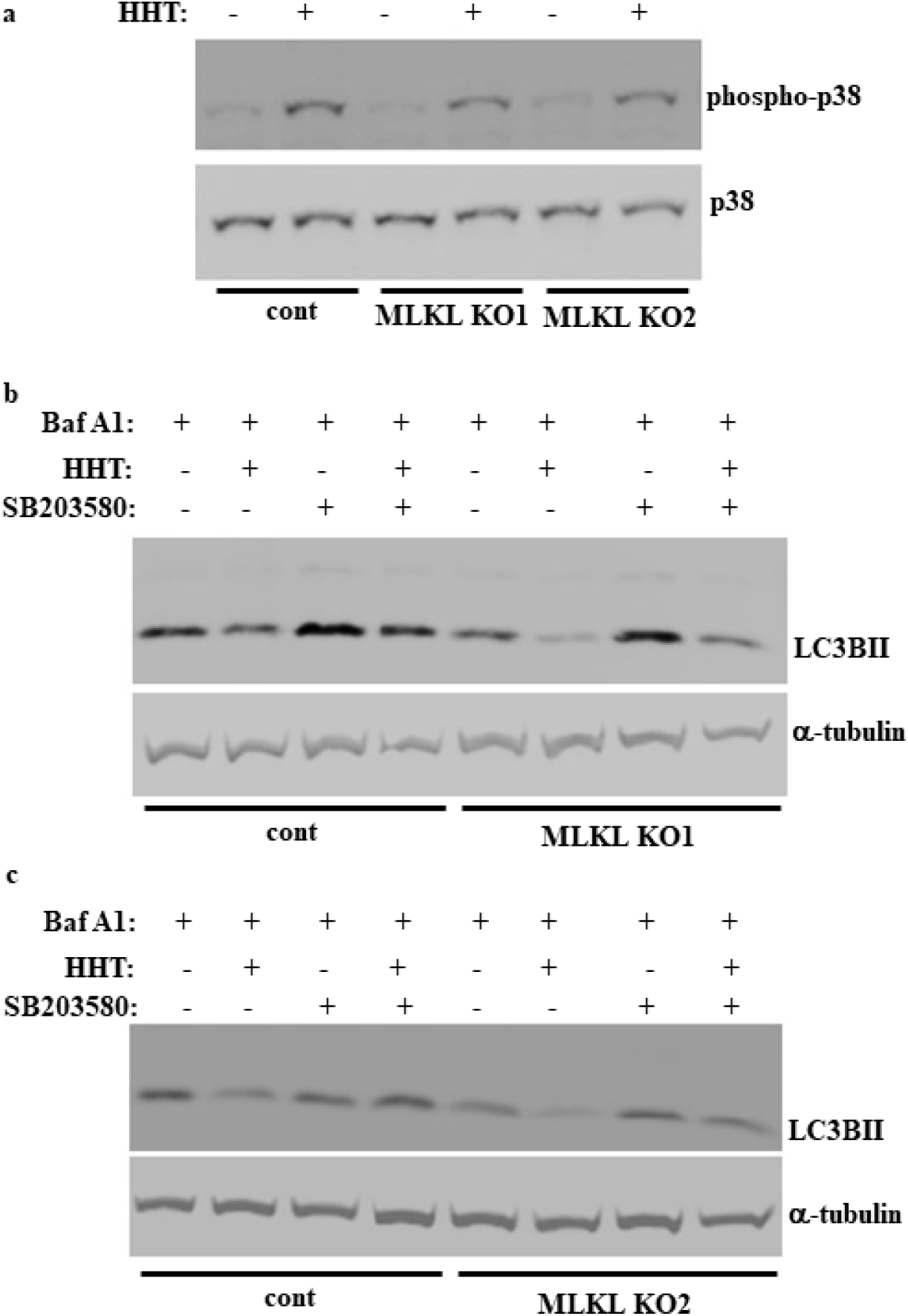
HHT contributes to autophagy inhibition in CRC cells by activating p38MAPK. Indicated cell lines were treated with DMSO (-) or 10 ng/ml HHT for 24h and/or 20 μM p38MAPK inhibitor SB203580 (SB203580) (+) in the presence of 50 nM bafilomycin A1 and assayed for p38MAPK and phospho-38MAPK (**a)** or LC3B (**b, c**) levels by western blotting. α-tubulin was used as a loading control.

### MLKL inactivation and HHT cooperate in killing CRC cells by parthanatos

Numerous forms of regulated cell death are presently known^2^, and we investigated which of these death forms is triggered by the combined inactivation of MLKL and HHT treatment. When using flow cytometry to analyze the distribution of cells between the phases of the cell cycle based on their DNA content, we noticed that treatment of the control MLKL-expressing DLD1 cells with HHT significantly increased the fraction of cells with the hypodiploid sub-G1 DNA content (Fig. 8a, b, g) and that percentage of the cells with the sub-G1 DNA is much higher in HHT-treated MLKL KO cells than in HHT-treated MLKL-expressing control cells (Fig. 8a-g). The presence of the sub-G1 population is a well-established indicator of chromosomal DNA fragmentation^49^, a sign of regulated cell death types, such as apoptosis and parthanatos^2^. When we cultured the MLKL KO DLD1 cells with HHT in the presence of zVAD-FMK, a widely used inhibitor of caspases which execute apoptosis^50^, this resulted in massive chromosomal DNA degradation regardless of whether the cells produced MLKL, and this degradation made the samples unsuitable for flow cytometry analysis (not shown). Notably, caspase inhibition was observed to trigger non-apoptotic cell death in multiple studies^51, 52^. Since we found that the ability of HHT to kill cells strongly depends on the presence of MLKL (Fig. 2), while zVAD-FMK killed HHT-treated cells, rather than protected them from death, and exerted this effect regardless of MLKL expression (not shown), we concluded that it is unlikely that HHT treatment and MLKL inactivation cooperate with each other in killing CRC cells by apoptosis.

**Fig. 8.**
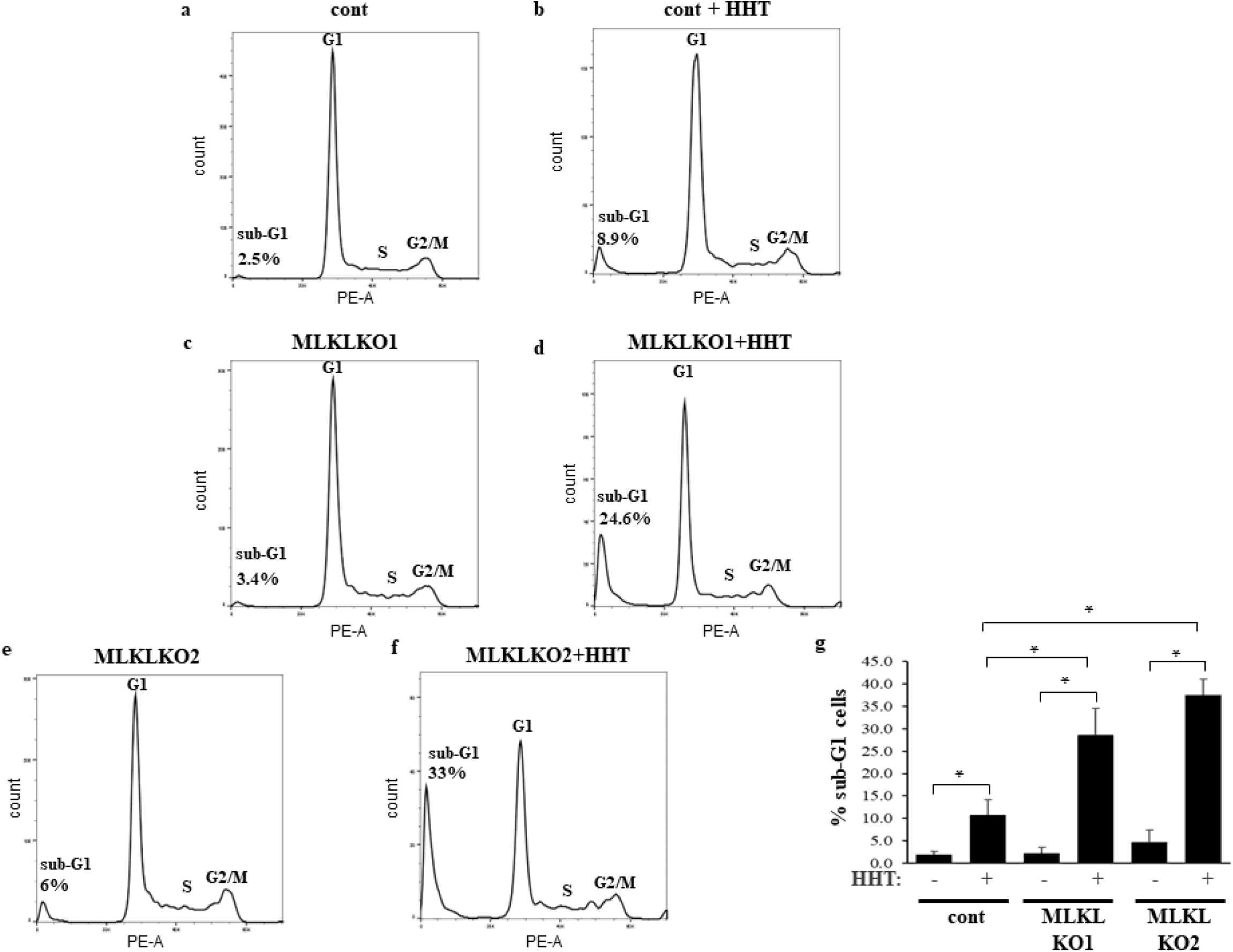
MLKL inactivation and HHT cooperate in triggering chromosomal DNA fragmentation in CRC cells. Indicated cell lines were treated with with 10 ng/ml HHT (HHT) or DMSO for 96h and analyzed for the cell cycle profile by flow cytometry. **a-f** Representative cell cycle profiles are shown. **g** The average % of the hypodiploid sub-G1 cells for the indicated treatments derived from three independent experiments plus SD is shown. *- p ˂0.05.

Another form of regulated cell death associated with DNA fragmentation is parthanatos^2^, programmed necrosis triggered by hyperactivation of poly(ADP-ribose) polymerase 1 (PARP1), an enzyme that drives the formation of poly(ADP-ribose) (PAR) in the cells^13^. When produced in excessive amounts, PAR binds a cell death-promoting factor AIF localized to the mitochondrial outer membrane^13^. This causes the release of AIF from the mitochondria. AIF then binds an endonuclease MIF, the AIF–MIF complex moves to the nucleus where it triggers chromosomal DNA degradation and cell death^13^.

We found that HHT-induced formation of the sub-G1 DNA content in the MLKL KO DLD1 cells is strongly inhibited by the PARP inhibitor olaparib^53^ (Fig. 9a-e). Moreover, treatment of these cells with HHT resulted in a significant increase in the multiple species of PAR polymers (Fig. 9f), an event that occurs in response to increased PARP activity and is one of the symptoms of parthanatos^54^. Thus, HHT and MLKL inactivation cooperate with each other in triggering CRC cell parthanatos. We further noticed that the autophagy-inducing Beclin1-derived cell-permeable peptide^35^ significantly reduced PAR species formation in HHT-treated cells (Fig. 9f), indicating that autophagy inhibition promotes parthanatos of HHT-treated MLKL-deficient CRC cells.

**Fig. 9.**
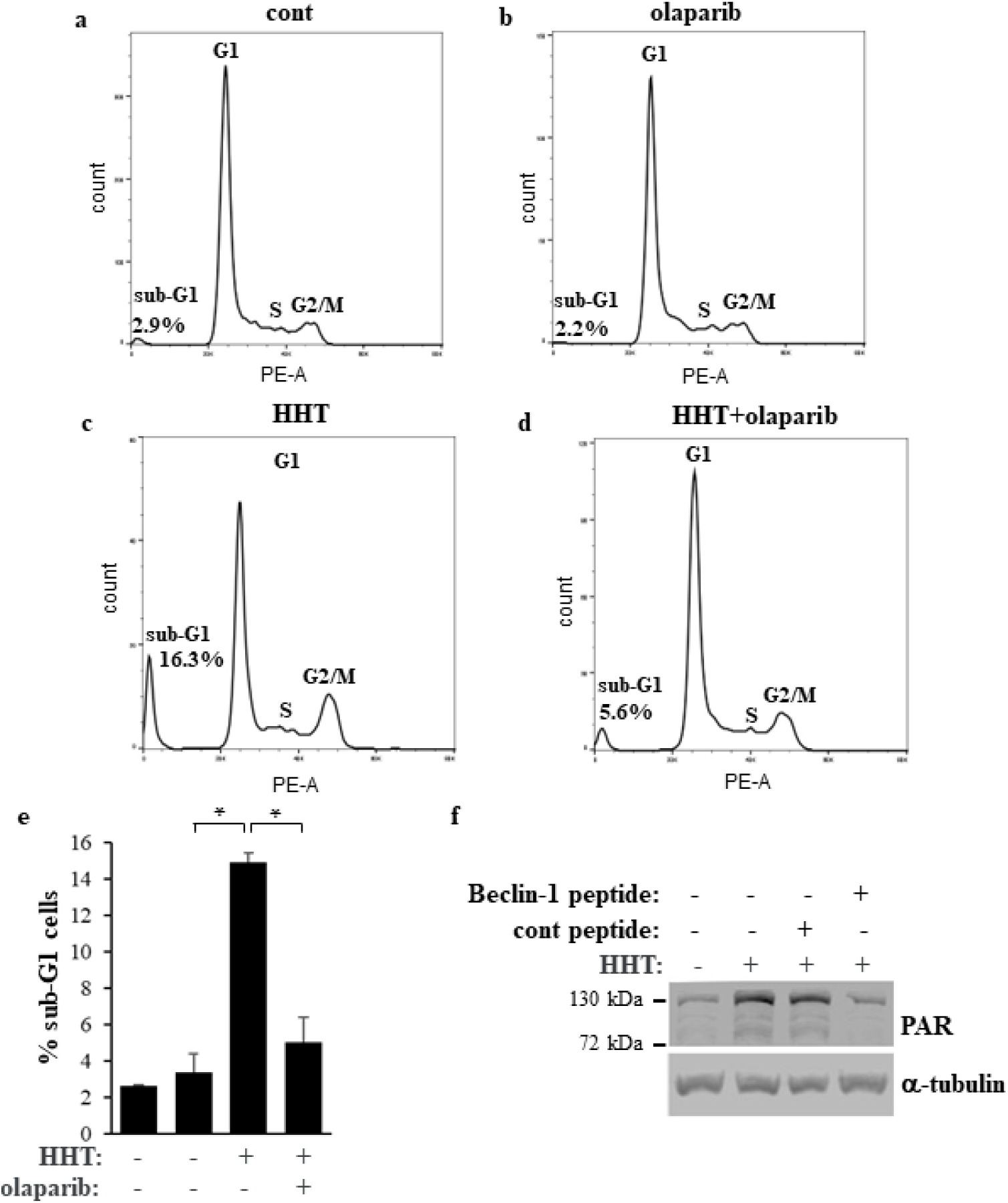
MLKL inactivation and HHT cooperate in triggering parthanatos of CRC cells. MLKL KO1 cells lines were treated with DMSO (cont), 10 ng/ml HHT (HHT), 50 nM olaparib (Olaparib), or both drugs (HHT+ Olaparib) at the indicated concentrations for 96h and analyzed for the cell cycle profile by flow cytometry. **a-d** Representative cell cycle profiles are shown. **e** The average % of the hypodiploid sub-G1 cells for the indicated treatments derived from three independent experiments plus SD is shown. *- p ˂0.05. (**f**) MLKL KO1 cells lines were treated with DMSO (-), 10 ng/ml HHT (+) in the absence (-) or in in the presence (+) of 12 μM of control (cont) or cell-permeable Beclin-1 (Beclin-1) peptide for 96h and analyzed for poly(ADP-ribose) (PAR) levels by western blotting. α-tubulin was used as a loading control.

### Low MLKL mRNA expression signifies increased CRC patient survival

Our data indicate that MLKL promotes CRC cell survival. If this is the case, then, conceivably, CRC cells producing low MLKL levels could be less viable and less deadly than those producing higher MLKL levels. To test whether MLKL expression correlates with CRC patient survival we used GEPIA, an interactive web server for analyzing the RNA sequencing expression data in various tumors derived from the TCGA and the GTEx projects^55^. We found that reduced levels of MLKL mRNA in CRC correlate with increased patient survival (**Fig. 10**). These data are consistent with the scenario whereby the lower the levels of MLKL expression in CRC cells are, the less viable the cells are, the less deadly respective cancers become, the longer patients survive.

**Fig. 10.**
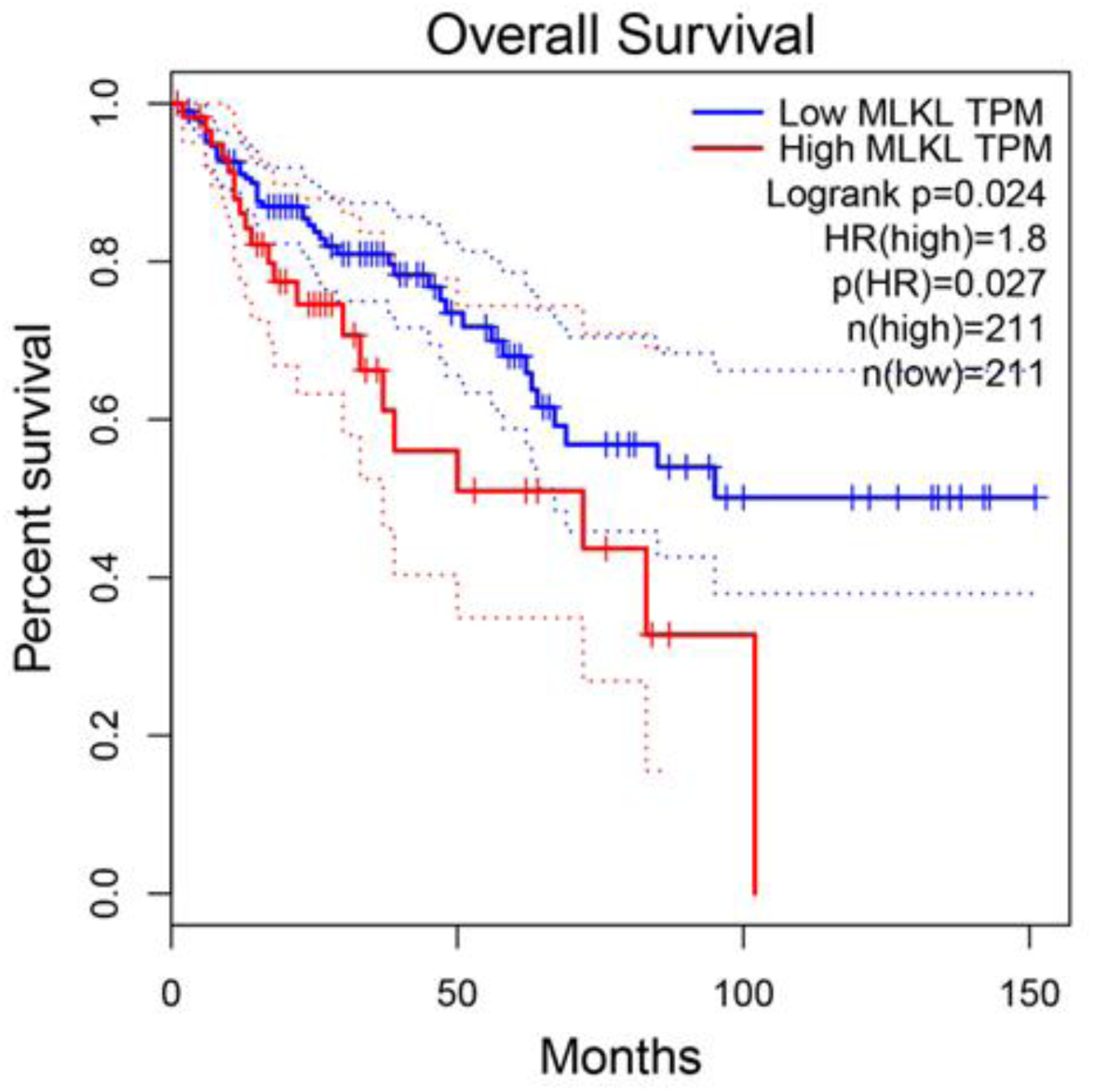
Low MLKL mRNA expression signifies increased CRC patient survival. Kaplan– Meier analysis-based estimation of probabilities of patients’ survival depending on the level of MLKL mRNA was performed.

In summary, we have demonstrated here that inactivation of MLKL, a protein well known for its ability to promote necroptosis, creates a therapeutic vulnerability in CRC cells in that they become more sensitive to treatment with HHT, a clinically approved anti-cancer drug. We have demonstrated that pharmacological or genetic MLKL inactivation, when combined with HHT, blocks the basal CRC cell autophagy and triggers their death by parthanatos. Moreover, we found that NSA, a pharmacological MLKL inhibitor, and HHT cooperate with each other in suppressing the ability of CRC cells to form tumors *in vivo*. Hence, pharmacological MLKL inhibition combined with homoharringtonine treatment represents a potential novel approach for CRC therapy.

## Discussion

A significant proportion of CRCs are incurable^1^, and novel CRC therapies are needed. We have demonstrated here that MLKL inactivation when combined with a clinically approved anti-cancer drug HHT represents an effective approach for killing CRC cells *in vitro* (Fig. 2) and suppressing growth of tumors formed by CRC cells *in vivo* (Fig. 3). Hence, combining MLKL inactivation with HHT treatment is a potential novel approach to CRC management.

We have also discovered here an unexpected role of MLKL in CRC cell survival. MLKL is well known to promote cell death in response to necroptosis-inducing stimuli^2^. We have found that MLKL can also support CRC cell survival (Fig. 1, 2, 8, 9). Notably, we found that reduced levels MLKL mRNA in CRC correlate with increased patient survival (Fig. 10). The indicated data are consistent with the scenario whereby the lower the levels of activated MLKL in the tumor cells are, the less viable the cells are, the less deadly respective cancers become.

We found that MLKL promotes CRC cell survival by supporting the basal autophagy in CRC cells (Fig. 5). We further discovered that MLKL inactivation and HHT treatment cooperate with each other in blocking such autophagy and killing CRC cells (Fig. 5). At the mechanistic level, our data are consistent with a scenario that MLKL mediates LC3B lipidation, one of the critical autophagy steps, in CRC cells. The ESCRT-I complex is known to promote autophagy^12^ but apparently, the ESCRT-I component VPS37A is not needed for driving LC3 lipidation while MLKL is expressed by the cells (Fig. 6e, f). However, in the absence of MLKL, the ability of the cells to induce LC3B lipidation is significantly reduced and seems to become almost entirely VPS37A-dependent (Fig. 6e, f). Notably, so far VPS37A was found to promote autophagy by supporting the autophagosomal membrane integrity and/or promoting the closure of this membrane during autophagy^11, 12^. To our knowledge the role of the ESCRT-I machinery in LC3 lipidation has never been demonstrated, which is not surprising since this role becomes apparent only in the absence of MLKL (Fig. 6e, f), a scenario that has not been explored until now. LC3 proteins are a members of the ATG8 protein family which also includes the GABARAP proteins^56^. Both LC3 and GBARAP proteins mediate autophagy via being lipidated. Notably VPS37A seems to participate in supporting the autophagosomal membrane integrity by physically binding LC3 and GABARAP proteins^11^. Hence, is conceivable that in certain circumstances, VPS37A can promote LC3 lipidation. Understanding how VPS37A controls such lipidation and how this process is regulated by MLKL is the subject of our ongoing studies.

We have demonstrated here that the ability of MLKL antagonists to suppress CRC cell autophagy is noticeably enhanced by HHT and that this effect is driven by HHT-induced p38MAPK activation (Fig. 7). HHT is an inhibitor of the activity of the ribosomes^6^ and such inhibition often results in ribotoxic stress response, a signaling cascade driven by p38MAPK^7^. Our data are consistent with these findings and with observations that p38MAPK is known to inhibit autophagy via multiple mechanisms^44, 45^.

Since both NSA and HHT inhibit autophagy, would it be possible to use either drug alone at a higher dose to suppress autophagy below the levels required for CRC cell survival in a clinical setting? We have found here that even complete MLKL loss has only a limited effect on the basal cell autophagy and cell survival (Fig. 1, 5) indicating that MLKL inactivation alone is insufficient for suppressing autophagy below cell survival-promoting level. Moreover, while HHT seems to show efficacy against CML^5^, the use of HHT as a single agent at non-toxic doses was ineffective against CRC in the clinic^57^, and a higher dose caused unacceptable toxicity in patients^58^. Thus, the use of MLKL antagonists in the clinic may render the non-toxic HHT doses sufficient for reducing autophagy below the levels required for CRC cell survival and thereby make this combination treatment efficacious against CRC.

We found that inhibition of autophagy caused by MLKL inactivation and HHT treatment kills CRC cells by parthanatos (Fig. 8, 9). Oxidative stress and excessive DNA damage are the known consequences of autophagy inhibition in cancer cells^59, 60^ and represent the established stimuli causing PARP hyperactivation and parthanatos^13^. Testing whether oxidative stress and/or DNA damage caused by autophagy inhibition trigger parthanatos in our model represents a promising direction for our future studies.

We demonstrated here that that NSA, a small molecule MLKL inhibitor^3^, strongly cooperates with HHT in suppressing growth of tumors formed by CRC cells in mice (Fig. 3). Thus, pharmacological MLKL inhibition combined with HHT treatment is a potential novel approach for CRC management. How could this approach improve the existing CRC treatments? It is known in this regard that PARP1, a major parthanatos inducer, is overexpressed in human tumor-derived colon cancer cells compared to normal intestinal epithelial cells and promotes colon cancer in mice^14^. Hence, high level and activity of PARP1 in CRC cells could make the cells vulnerable to PARP1-hyperactivating treatments, e.g. those based on MLKL inhibition combined with the use of HHT, and thereby create a therapeutic window for CRC treatment with these drugs. Moreover, MLKL gene KO is non-toxic to mice^4^, while HHT is already used for chronic myelogenous leukemia and is typically well tolerated by patients^5^. Therefore, adding this novel combination treatment to current CRC therapies could improve their efficacy, and PARP1 and/or other PARP isoforms could be explored as biomarkers of CRC sensitivity to this combination treatment.

## Methods

### Cell culture

Cell lines DLD1, HCT116, LS180, SW480 and 293T were cultured in DMEM (GIBCO) and 10% Fetal Bovine Serum (FBS) (Sigma), 100 U/ml penicillin (GIBCO), 100 μg/ml streptomycin (GIBCO), 0.29 mg/ml L-glutamine (GIBCO). Lack of mycoplasma contamination in all cells was established as published^61^.

### Reagents

The following reagents were from Cell Signaling Technology, Danvers, MA. Bafilomycin A1 (catalogue# 54645S), Z-VAD(OMe)-FMK (catalogue# 60332S), Olaparib (catalogue# 93852S), SB203580 (catalogue# 5633S). The following other reagents were used. Vemurafenib, Santa Cruz Biotechnology, Dallas TX, USA (catalogue# CAS 918504-65-1), selumetinib, Cedarlane Labs, Burlington ON, Canada (catalogue# ENZ-CHM184-0050), pictilisib, Cedarlane Labs (catalogue# HY-50094-10MG), lapatinib Selleckchem, Houston, TX, USA (catalogue# S1028), Lipofectamine™ 3000 Transfection Reagent, Thermo Fisher Scientific Waltham, MA, USA (catalogue# L3000008), Tat-BECN1 peptide, Selleckchem Houston, TX, USA (catalogue# S8595), scrambled peptide, MedChemExpress, Monmouth Junction, NJ, USA (catalogue# HY-P4106).

### Antibodies

The following antibodies used for western blotting were from Cell Signaling Technology, Danvers, MA, USA. Anti-LC3B (D11) (catalogue# 3868S), anti-DR4 (D9S1R) (catalogue# 42533S) anti-DR5 (D4E9) (catalogue# 8074S), anti-MLKL (D2I6N) (catalogue# 14993S), anti-alpha-tubulin (DM1A) (catalogue# 3873S), anti-poly/Mono-ADP Ribose (D9P7Z) (catalogue# 89190S), anti-p38 (catalogue# 9212), phospho-p38 (catalogue# 4511). Anti-VPS37A was from Proteintech, Rosemont, IL, USA (catalogue# 11870-1-AP).

### MLKL gene knockout by CRISPR

sgRNA-encoding cDNAs targeting exon 2 (TTGAAGCATATTATCACCCT) or exon 9 (TTAGCTTTGGAATCGTCCTC) of the MLKL gene were generated by use of CHOPCHOP web tool version 3^62^ and cloned into pX459 expression vector. DLD-1 cells were transfected for 1 day as described with 2ug of pX459 encoding each sgRNA cloned into pX459^63^, the cells were treated with 3μg/ml of puromycin for 2 days, and clonal cell lines were isolated by limiting dilution and analyzed for MLKL expression by western blotting. A clone that still expressed MLKL was chosen as a control clone, and clones that did not show MLKL expression in the case of each sgRNA were selected as MLKL-deficient clones.

### RNA interference

Small interfering (si)RNAs (Thermo Fisher Scientific, Waltham, MA, USA) were used as described^64^. Silencer™ Select Negative Control No. 1 siRNA (cat # 4390843) served as a control. The sequences of VPS37A siRNA1 and 2, respectively, were CGACAAAGAUGACUUAGUATT (cat #s44037) and CGACAUCACUUAAUGGAUATT (cat# s44038).

### Drug treatment for the colony formation assay

In the case of treatment of cells with NSA, HHT or other drugs described in the study, cells were plated on day 1, treated with NSA on day 2, with the second drug, on day 3, and the medium was changed to drug-free medium on day 7. In the case of MLKL KO clones, cells were plated on day 1, treated with a drug on day 2, and the medium was changed to drug-free medium on day 5.

### Tumor xenografts

Animal studies were approved by Dalhousie University Committee on Laboratory Animals. 6-week-old Nu/Nu Nude mice (Charles River Canada, Saint-Constant, QC) were allowed to acclimatize for 2 weeks. 8 × 10^6^ cells were harvested by trypsin treatment, washed thrice in ice-cold PBS and injected in the muse flank. When tumor volumes reached the average of 80 mm^3^, the mice were injected intraperitoneally (IP) with either DMSO or 4.2mg/kg NSA or 1mg/kg HHT or both. NSA was injected daily for 7 days and HHT, daily for 5 days, followed by a 2-day break followed by 5 more days of daily injections. Tumor volumes were measured as published^23^.

### Hypodiploid sub-G1 DNA content analysis

20,000 cells/well were seeded in a 6-well plate, harvested after drug treatment using 0.25% trypsin at 37°C for 5 minutes, followed by trypsin neutralization with equal volume of the respective serum-containing medium. Cells were pelleted at 500 g for 5 min, washed with 2 ml phosphate-buffered saline (PBS), pelleted at 500 g at 4°C for 5 min, fixed with 5 ml of 70% ethanol overnight at 4°C, pelleted at 900 x g for 10 min, washed with 3ml PBS, pelleted at 900 g at 4°C for 5 min, resuspended in 500 μl of PBS containing 100 μg/mL RNase A, incubated for 30 min at room temperature. 200 µl of propidium iodide was then added to the cells. The cells were analyzed by flow cytometry using BD FACS Celesta instrument, and the data were processed using Flowjo software.

### Assessment of probability of patient’s overall survival

The analysis was carried out by use of GEPIA web tool^55^. The cut off for low and high MLKL mRNA expression levels was 78% and 22%, respectively.

Western blotting^61^, detection of clonogenic cell survival^65^ and detection of the LC3 puncta^66^ were performed as published^65^.

### Statistical Analysis

Statistical analysis of the data in Fig. 1a and 9e was performed by the two-sided chi-square test for goodness-of-fit, and statistical analysis of all other data, by the two-sided Student’s t-test.

## Acknowledgements

This study was supported by grants from The Cancer Research Society and Canadian Institutes of Health Research (KVR, principal investigator on both grants). XL and SC were recipients of Cancer Research Training Program postdoctoral fellowships via Beatrice Hunter Cancer Research Institute. SC and PJ were recipients of the IWK Health Sciences Centre postdoctoral fellowships.

## Contributions

PJ, BHY, SC and XL performed the experiments. KVR conceived and supervised the study as well as wrote the initial draft of the manuscript.

## Conflict of interest

We have no conflicts to declare.

## Ethics approval

Animal studies were approved by Dalhousie University Committee on Laboratory Animals.

## Data availability

All data reported in this study will be shared by us upon request.

## Notes

### Competing Interest Statement

The authors have declared no competing interest.

